# Electrophysiological analyses of human dorsal root ganglia and human induced pluripotent stem cell-derived sensory neurons from male and female donors

**DOI:** 10.1101/2023.11.03.565343

**Authors:** Nesia A. Zurek, Reza Ehsanian, Aleyah E. Goins, Ian M. Adams, Timothy Petersen, Sachin Goyal, Mark Shilling, Karin N. Westlund, Sascha R.A. Alles

## Abstract

Human induced pluripotent stem cell-derived sensory neurons (hiPSC-SNs) and human dorsal root ganglia (hDRG) neurons are popular tools in the field of pain research; however, few groups make use of both approaches. For screening and analgesic validation purposes, important characterizations can be determined of the similarities and differences between hDRG and hiPSC-SNs. This study focuses specifically on electrophysiology properties of hDRG in comparison to hiPSC-SNs. We also compared hDRG and hiPSC-SNs from both male and female donors to evaluate potential sex differences. We recorded neuronal size, rheobase, resting membrane potential, input resistance, and action potential waveform properties from 83 hiPSCs-SNs (2 donors) and 108 hDRG neurons (9 donors). We observed several statistically significant electrophysiological differences between hDRG and hiPSC-SNs, such as size, rheobase, input resistance, and several actional potential (AP) waveform properties. Correlation analysis also revealed many properties that were positively or negatively correlated, some of which were differentially correlated between hDRG and hiPSC-SNs. This study shows several differences between hDRG and hiPSC-SNs and allows better understanding of the advantages and disadvantages of both for use in pain research. We hope this study will be a valuable resource for pain researchers considering the use of these human *in vitro* systems for mechanistic studies and/or drug development projects.

## INTRODUCTION

Human induced pluripotent stem cell-derived sensory neurons (hiPSC-SNs) and human dorsal root ganglia neurons (hDRG-N) are popular tools in the field of pain research; however, few groups make use of both approaches. We and other groups have made use of these *in vitro* tools for the purposes of validation of novel analgesics and targets using whole-cell patch-clamp electrophysiology[5,7,10,32,33,40]. Since differences in source, genetic makeup, developmental stage, and functional characteristics can influence recorded data, our objective is to compare the electrophysiological properties of recorded hiPSC-SN and hDRG-N under controlled conditions.

Utilizing donor-derived hDRG-N for live cell recordings can pose challenges due to the complexities of acquiring fresh tissue. Furthermore, hDRG primary cultures contain multiple cell types including nociceptors, satellite glia, and immune cells. Additionally, the heterogeneity of donor demographics such as age, sex, prior opioid use, and disease state can introduce variability in neuronal excitability and responsiveness to analgesics. These characteristics of hDRG-N make homogenous hiPSC-SNs an attractive potential tool for electrophysiology experiments. Since hiPSC-SNs are more homogenous, they offer lower experimental variability and greater power to observe changes in electrophysiological properties[6]. However, hiPSC-SNs lack the factors that are present within the microenvironment of the dorsal root ganglia and may not be a true reflection of *in vivo* responses. The genetic makeup of hiPSC-SNs may also not perfectly replicate the genetic characteristics of the person from whom they were derived. More importantly since hDRG-N are taken from different donors they reflect the genetic make up of the specific donor without the issues of molecular manipulation of hiPSC-SNs. Furthermore, the differentiation process of hiPSC-SNs can be controlled, but it may not perfectly replicate the developmental stages and maturation of naturally occurring sensory neurons. hDRG-N are mature neurons in their natural state, so they represent neurons at a specific stage of development. The functional characteristics of hiPSC-SNs may exhibit functional differences in terms of their electrophysiological properties, responsiveness to stimuli, and other characteristics; the differentiation process and culture conditions can influence their properties. In comparison, hDRG-N are known to be fully functional sensory neurons and may more accurately represent the characteristics and responses of native sensory neurons in the human body.

For screening and drug validation purposes, important functional and genetic characteristics of hDRG-N and hiPSC-SNs can be determined using patch-clamp electrophysiology, mRNA, and protein expression analyses[5,44]. This paper is dedicated to examining the functional attributes of hDRG-N by analyzing their electrophysiological properties in comparison to those of hiPSC-SNs. Additionally, we conducted comparisons between hDRG-N and hiPSC-SNs obtained from male and female donors to assess the potential presence of sex-based differences.

We recorded intrinsic properties such as neuronal size, rheobase, resting membrane potential, input resistance, as well as action potential (AP) waveform properties from 83 hiPSCs-SNs (2 donors) and 108 hDRG-N (9 donors). We observed several statistically significant electrophysiological differences between hDRG-N and hiPSC-SNs. This study reveals numerous distinctions between hDRG-N and hiPSC-SNs, facilitating a more comprehensive understanding of their respective merits and limitations for application in pain research.

## METHODS

### Human dorsal root ganglia (hDRG) Culture

hDRG were obtained from recently deceased organ donors (Figure 1A). All procedures were performed at UNM Hospital in coordination with New Mexico Donor Services after obtaining appropriate consent. Study activities were approved by the Human Research Review Committee at the University of New Mexico Health Sciences Center; approval numbers #21-412 or #23-205. Cultures of hDRG-N (Figure 1B) were prepared as described previously[40] and cultured for up to 11 days *in vitro* (DIV). Electrophysiological recordings took place between DIV 3-11. There were 108 hDRG-N (59 male and 49 female cells) selected that were naïve or vehicle (media or water) controls. The demographics of the donors are shown in Figure 1C.

**Figure 1:**
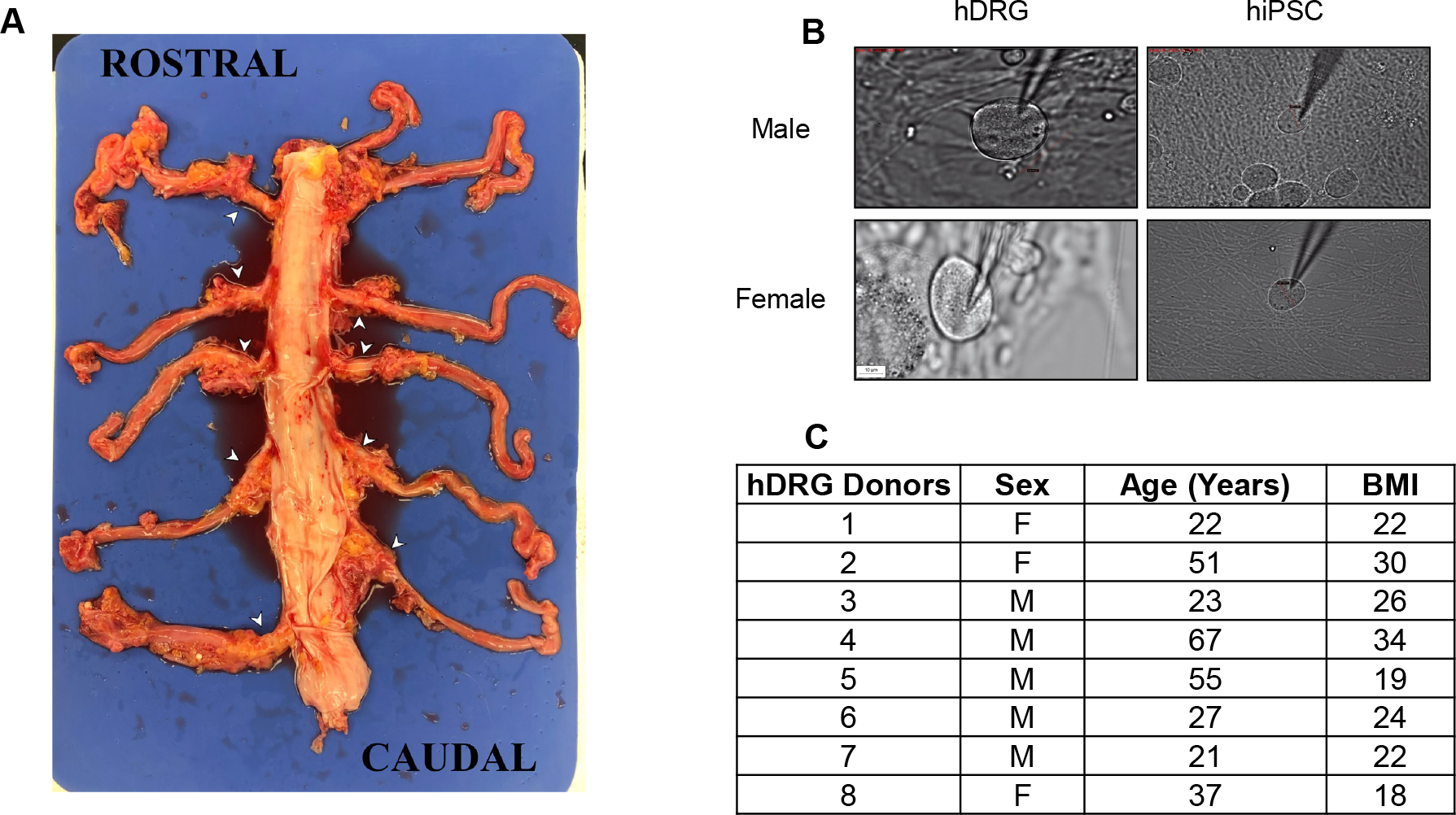
Culture of hDRG and hiPSC-SNs. **A**. Representative donor spinal cord and DRG (white arrows) used for culture of hDRG. **B**. Cell images of male and female hDRG and hiPSC-SNs. **C**. Donor demographics for hDRG used in this study.

### Human induced pluripotent stem cell-derived sensory neurons (hiPSC-SNs)

hiPSC-SNs were obtained from Anatomic Corp (Minneapolis, MN) and produced from female subject ANAT001 or a male subject. Cells were grown in culture as described previously[10] and were maintained in culture for up to 34 DIV (Figure 1B). Electrophysiological recordings took place between DIV 7-34 as performed previously[10]. Considered in these data are 83 naïve, untreated control cells (either media or water), 50 from males and 33 from females.

### Whole-cell patch-clamp electrophysiology

Whole cell patch-clamp electrophysiology was performed as previously described [10,11]. Recordings were done at room temperature, with the recording chamber perfused with artificial cerebrospinal fluid (aCSF) containing 113 mM NaCl, 3 mM KCl, 25 mM NaHCO3, 1 mM NaH_2_PO_4_, 2 mM CaCl_2_, 2 mM MgCl_2_, and 11 mM D-glucose bubbled with 95% O_2_/5% CO_2._ Neurons were identified with differential interference contrast optics connected to an IR-2000 digital camera (Dage MTI, MI) or an Olympus digital camera. Size was measured using Dage MTI (Indiana City, MI) camera software or ImageJ (NIH, Bethesda, MD). Current-clamp recordings were performed using a Multiclamp 700B (Molecular Devices, San Jose, CA). Signals were filtered at 5 kHz, acquired at 50 kHz using a Digidata 1550B converter (Molecular Devices, San Jose, CA), and recorded using Clampex 11 software (Molecular Devices, San Jose, CA). Patch pipettes were made with a Zeitz puller (Werner Zeitz, Martinsreid, Germany) from borosilicate thick glass (GC150F, Sutter Instruments). Electrode resistance was 3-7 MΩ. Bridge balance was applied for all recordings. Intracellular solution contained 120 mM K-gluconate, 11 mM KCl, 1 mM CaCl_2_, 2 mM MgCl_2,_ 10 mM HEPES, 11 mM EGTA, 4 mM Mg-ATP. Cell capacitance was calculated using the whole-cell capacitance compensation circuit in Multiclamp 700B. Exclusion criteria consisted of cells that did not fire APs, had a resting membrane potential (RMP) greater than −35 mV, or an access resistance greater than 15 MΩ.

Analysis was performed in Easy Electrophysiology v.2.5.1 (London, UK) and Clampfit 11.2 (Molecular Devices, San Jose, CA). All statistical analysis was performed using GraphPad Prism v10.0.2 (Boston, MA). Error bars denote mean ± SEM unless otherwise specified. Analysis methods are detailed in the methods section and figure texts.

### Electrophysiology intrinsic properties analysis methods

For current clamp recordings, a 500 ms current increasing stepwise from −100 pA in increments of 10 pA was typically the first protocol run on a patched cell. Rebound firing was noted if any APs occurred at the offset of the hyperpolarizing current injections. RMP was calculated from the average of 10 sweeps minimum. Input resistance was determined from 10 hyperpolarizing traces and the measured magnitude of the voltage deflection. Steady state voltage (V_SS_) was measured over approximately the last 100-200 ms of the current injection. The current-voltage relationship linear regression generated by Easy Electrophysiology was visually inspected along with the traces to identify disrupted recordings.

Rheobase is measured at the current injection that elicits firing, not considering any rebound firing or spontaneous, non-evoked activity, thus the minimum possible value is 10 pA. Cells were continually stimulated stepwise until they were observed to reach rheobase and then further activated until they reached inactivation (the point where the cell was no longer firing or started having fewer APs after the maximum number of APs), up to 4 nA. Cells that fired more than one AP during any current injection step at rheobase or higher were considered to be multi-firing. APs from spontaneous activity were excluded in determining multi-firing cells.

Sag was calculated in Easy Electrophysiology software at a 500 ms hyperpolarizing −100 pA current injection at the minimum voltage (V_sag_) minus the steady state voltage during the last 100ms of current injection (V_SS_). The voltage change from the resting membrane potential (RMP) of the sag voltage (ΔV_sag_) and the voltage change from RMP of the steady state voltage (ΔV_SS_) were used to calculate the sag ratio using the following equation.

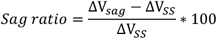

First spike latency was calculated in Easy Electrophysiology software by measuring the time from the start of current injection to the time of the first action potential at rheobase.

Statistics for all intrinsic properties were performed in GraphPad Prism v10.0.2 using an ordinary one-way ANOVA and Šídák’s multiple comparisons test.

### Frequency-current (f-I) relationship plots

The f-I plots were generated by considering only multi-firing cells (72 hDRG-N and 58 hiPSC-SN). Number of APs generated at current above rheobase was plotted and analyzed in GraphPad Prism 10 using mixed-effects analysis (REML) and Šídák’s multiple comparisons test at each current level. Inactivation current of multi-firing cells was defined as the current step at which cells started having fewer APs after the max number of APs. Statistics for inactivation threshold were performed in GraphPad Prism v10.0.2 using an ordinary one-way ANOVA and Šídák’s multiple comparisons test.

### Action potential waveform and phase plot analysis

The AP generated at rheobase was used for AP waveform and phase plot analysis. In Easy Electrophysiology AP kinetics analysis was used to generate waveform and phase plot data. The AP peak was defined as the highest mV of the AP. For thresholding, Method II[34] was used with constraints looking up to 4 ms before the AP peak and visually verified. The AP amplitude was defined as the change in mV from the threshold to the AP peak. AP rise and decay time was done using 10-90% cutoffs. The after hyperpolarization (AHP) was defined as the difference between the threshold and the lowest value of the AP hyperpolarization voltage value. The half-width time was automatically calculated by Easy Electrophysiology using those constraints. Statistics were performed in GraphPad Prism v10.0.2 using an ordinary one-way ANOVA and Šídák’s multiple comparisons test.

Phase plots were generated in Easy Electrophysiology software by looking 20 ms before the AP peak and 50 ms after the AP peak to generate a full phase plot for all APs. Phase plots were used to determine AP shoulder prevalence by looking at the decay part of the curve[7]. If the slope (dVm) increased for the entirety of the curve or increased, plateaued, and then increased again the curve was defined as having no AP shoulder. If the dVm had a distinct hump of the decay curve where it decreased, then increased, then decreased again the AP was defined as having an AP shoulder. For average phase plots the Vm across the population was averaged while the individual dVm values were used to generate the standard error of the mean (SEM) calculated in GraphPad Prism v10.0.2. All data was plotted in GraphPad Prism v10.0.2.

### Correlation Plots

Pairwise Pearson correlation matrices were generated for RMP, input resistance, rheobase, first-spike latency, sag ratio, maximum count of APs, half-width, and AHP using JMP 9.0.0 software (SAS Institute Inc., Cary NC, USA). The matrices were produced for pooled-sex samples of hDRG-N and hiPSC-SN, as well as separate male and female subsets of each.

## RESULTS

### Intrinsic properties of hDRG-N versus hiPSC-SNs

Discernible variations in intrinsic characteristics between hDRG-N and hiPSC-SNs were identified across various properties. Cell diameter measured during clamp recording with an IR-2000 camera (Figure 1A-D) showed the hDRG-N are significantly larger in diameter, with mean and SEM of 45.07 ± 0.93 µm, compared to hiPSC-SN, 21.65 ± 0.405 µm, p < 0.0001 (Figure 2A). The resting membrane potential (RMP) between hDRG-N (−54.92 ± 0.71 mV) and hiPSC-SNs (−56.06 ± 0.908 mV) was not significantly different (Figure 2B). A significant difference in rheobase was detected between hDRG-N and the hiPSC-SNs (592.7 ± 63.02 pA versus 74.22 ± 6.756 pA, p < 0.0001) (Figure 2C). R_in_ (input resistance) was compared showing no significant difference between hDRG-N and hiPSC-SNs (248.2 ± 31.12 MΩ versus 339.2 ± 12.55 MΩ) (Figure 2D). Capacitance was significantly different between the hDRG-N and hiPSC-SN, (104.0 ± 26 ± pF vs 23.06 ± 1.113 pF, p < 0.0001). The sag percentage at −100 pA ratio was significantly different between the hDRG-N and hiPSC-SNs (23.87 ± 1.562 vs 17.10 ± 1.104, p = 0.0027) (Figure 2F). The first spike latency (FSL) was significantly different between hDRG-N and hiPSC-SNs with the hDRG-N firing later at 96.16 ± 113.45 ms and hiPSC-SNs firing sooner at 46.06 ± 5.028 ms after the start of current injection, p = 0.0089. The hiPSC-SNs intrinsic properties were more consistent and had smaller SEM across almost all properties including cell diameter, rheobase, R_in_, capacitance, and sag ratio than hDRG-N.

**Figure 2.**
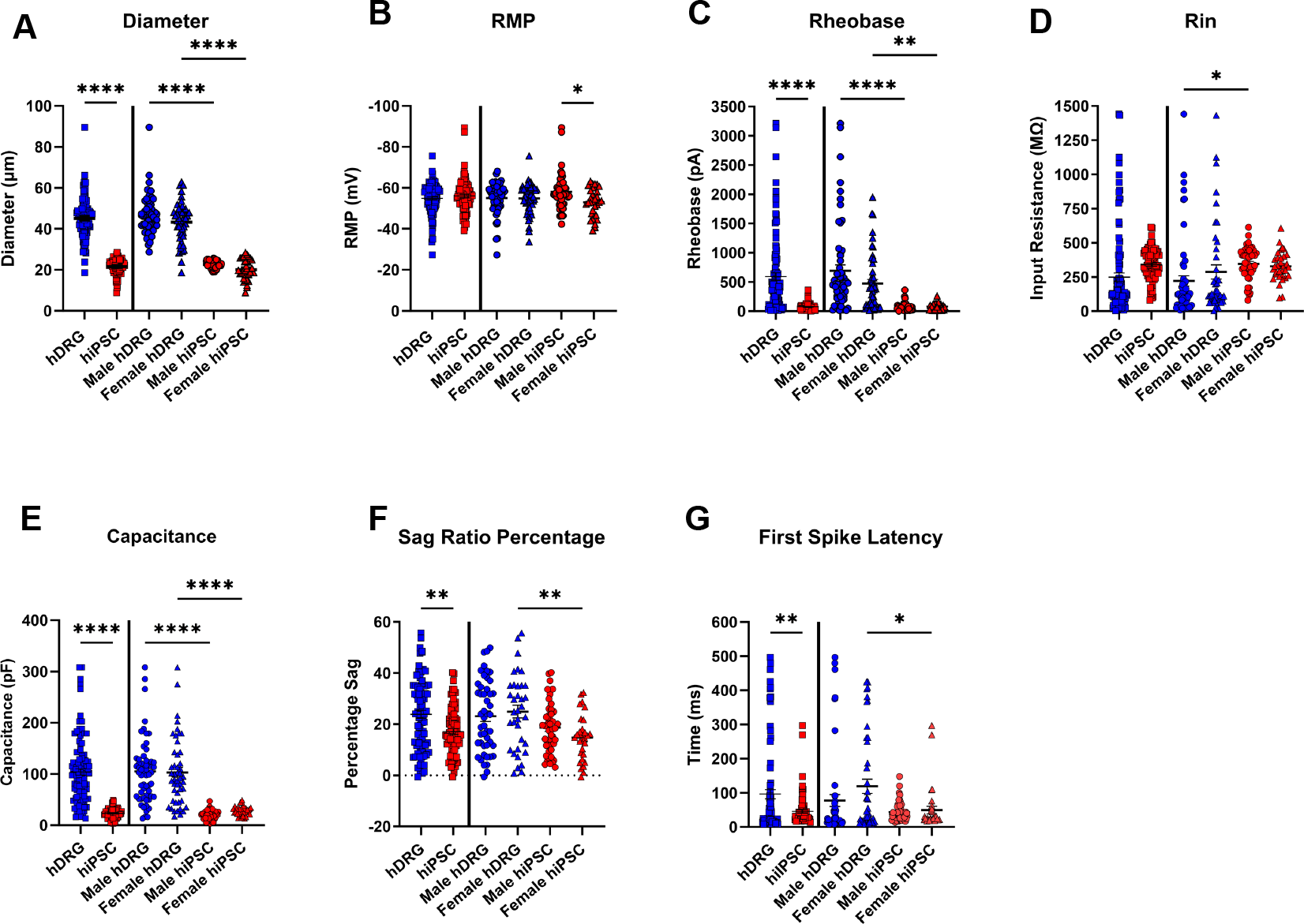
Intrinsic electrophysiological differences were observed between hDRG cells (n=108) and hiPSC-SNs (n=83). **(A)** Cell diameter was confirmed during patch clamp recording with an IR-2000 camera. **(B)** resting membrane potential (RMP), **(C)** rheobase, and **(D)** input resistance were recorded in current clamp mode. (**E)** capacitance (pF) was measured by whole-cell capacitance. **(F)** sag ratio. **(G)** first Spike Latency. Ordinary one-way ANOVA was used to determine statistical significance between hDRG cells (n=59 from males, 49 from females) and hiPSC-SNs (n=50 from male, 33 from female). Šídák’s multiple comparisons test confirmed significant sex differences were detected in each intrinsic property. *p=.01, **p<.01, ***p<.001, ****p<.0001. Error bars represent mean and SEM.

### Excitability of hDRG-N compared to hiPSC-SNs

Excitability of the hDRG-N and hiPSC-SNs were analyzed by performing stepwise current injection up to at least 150 pA above rheobase (Figure 3A). APs that occurred during the 500 ms current injection and not any rebound or spontaneous APs were used in determining multi-firing cells and for measuring the number of APs during each current clamp step (Figure 3A). The hiPSC-SNs had more APs at every current level up to at least 150 pA over rheobase (Figure 3B) when compared to the hDRG-N, p < 0.0001 at 150 pA. These data show that hiPSC-SNs are in general more excitable than hDRG-N.

**Figure 3.**
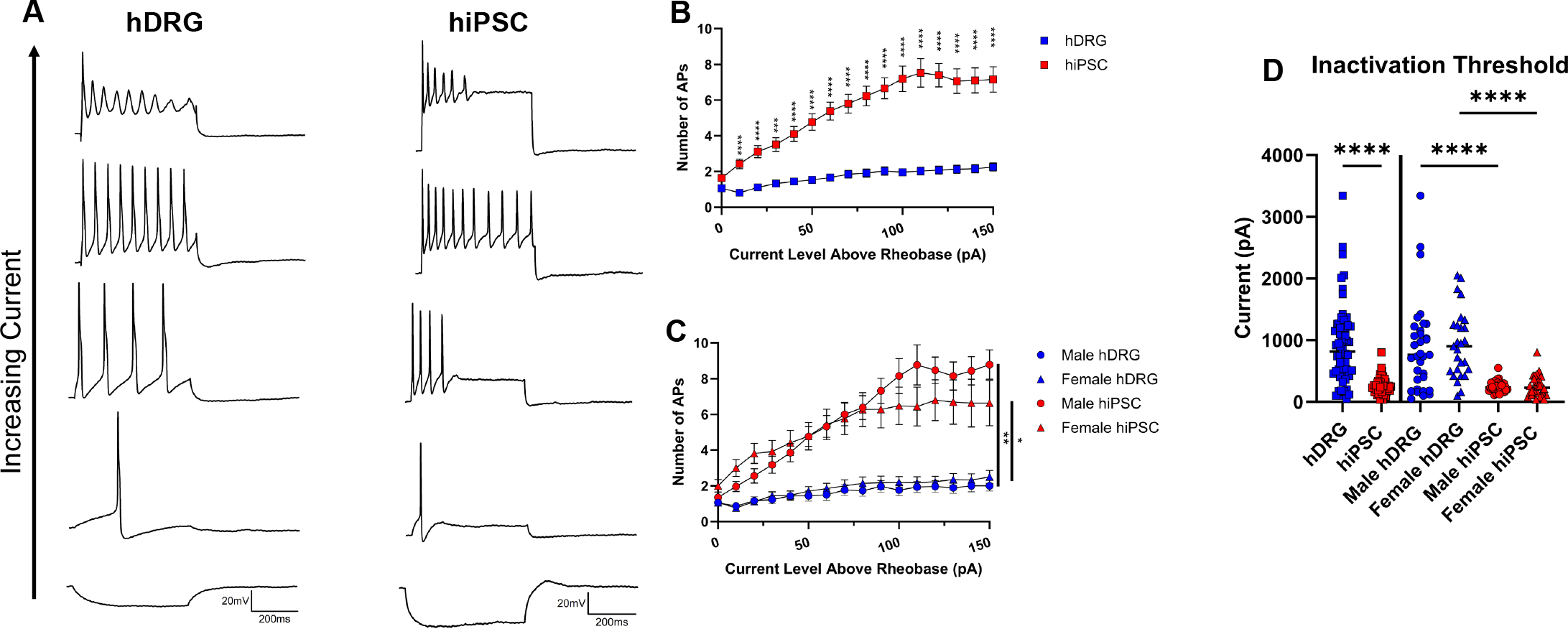
Frequency measured as a function of injected current level above rheobase. Representative traces of **(A)** hDRG and hiPSC-SNs examples of firing at −100pA, rheobase, multi-firing, and inactivation. **(B)** hDRG (n=72 cells) and hiPSC-SNs (n=58 cells) had significant differences of excitability in multi-firing cells except at baseline rheobase. Repeated measures ANOVA for the linear mixed-effects model (REML) was performed with Šídák methods applied to evaluate multiple comparisons between the mean of action potential (APs) frequency over an injected current level above rheobase (pA) for hDRG and hiPSC-SNs multi-firing cells. Inclusion criteria defined as firing 2 or more action potentials during a step protocol in current clamp mode until inactivation or at least 150pA above rheobase. Greenhouse-Geisser correction applied for unmet sphericity assumption (epsilon<0.75). ***p=.001, ****p<.0001. **(C)** Male hDRG (n=35) and female hiPSC-SNs (n=37) versus male hiPSC-SNs (n=33) and female hiPSC-SNs (n=25), respectively. Error bars represent SEM. **(D)** inactivation threshold for multi-firing cells defined as when cells started having less APs after reaching max APs. Ordinary one-way ANOVA was used to determine statistical significance. ****p<.0001. Error bars represent mean and SEM.

Inactivation threshold of each group, defined as the current level over rheobase when the cells started having fewer APs or stopped firing, was analyzed (Figure 3D). The hDRG-N continued to fire well above rheobase (909.8 ± 84.45 pA), compared to hiPSC-SN (249.3 ± 17.6 pA), p <0.0001. These data show that hDRG-N can continue to fire well over rheobase while hiPSC-SN inactivate at much lower current levels.

### Phenotypic firing patterns of hDRG compared to hiPSC-SNs

The prevalence of multi-firing cells was not statistically significant between hiPSC-SNs and hDRG-N, [71% (58/82) vs 66% (72/108)] (Figure 4B, G). Rebound firing was defined as cells that had an AP after the 500 ms current injection. The hiPSC-SNs, 47% (44/83), were much more likely to have rebound firing than hDRG-N, only 4% (5/108), p < 0.0001 (Figure 4C, G). Spontaneous activity was determined by analyzing APs during 30 s gap-free current clamp steps where there was either no current applied or enough current to hold the cell’s membrane potential at −45mV. When combined, hDRG-N, 34% (31/91), were more likely than hiPSC-SN, 19% (14/75), to have spontaneous activity, p = 0.0244 (Figure 4D, G). hDRG-N were also more likely than hiPSC-SNs to display delayed firing at rheobase (over 100 ms after current input started), with 33% (36/108) versus 7% (6/83) respectively, p < 0.0001 (Figure 4E, G). There was a significant percentage difference in cells with visible sag (more than 10% sag percentage) with 53% (53/100) of hDRG-N compared to 73% (58/79) of hiPSC-SNs displaying visible sag, p = 0.0052 (Figure 4F, G).

**Figure 4:**
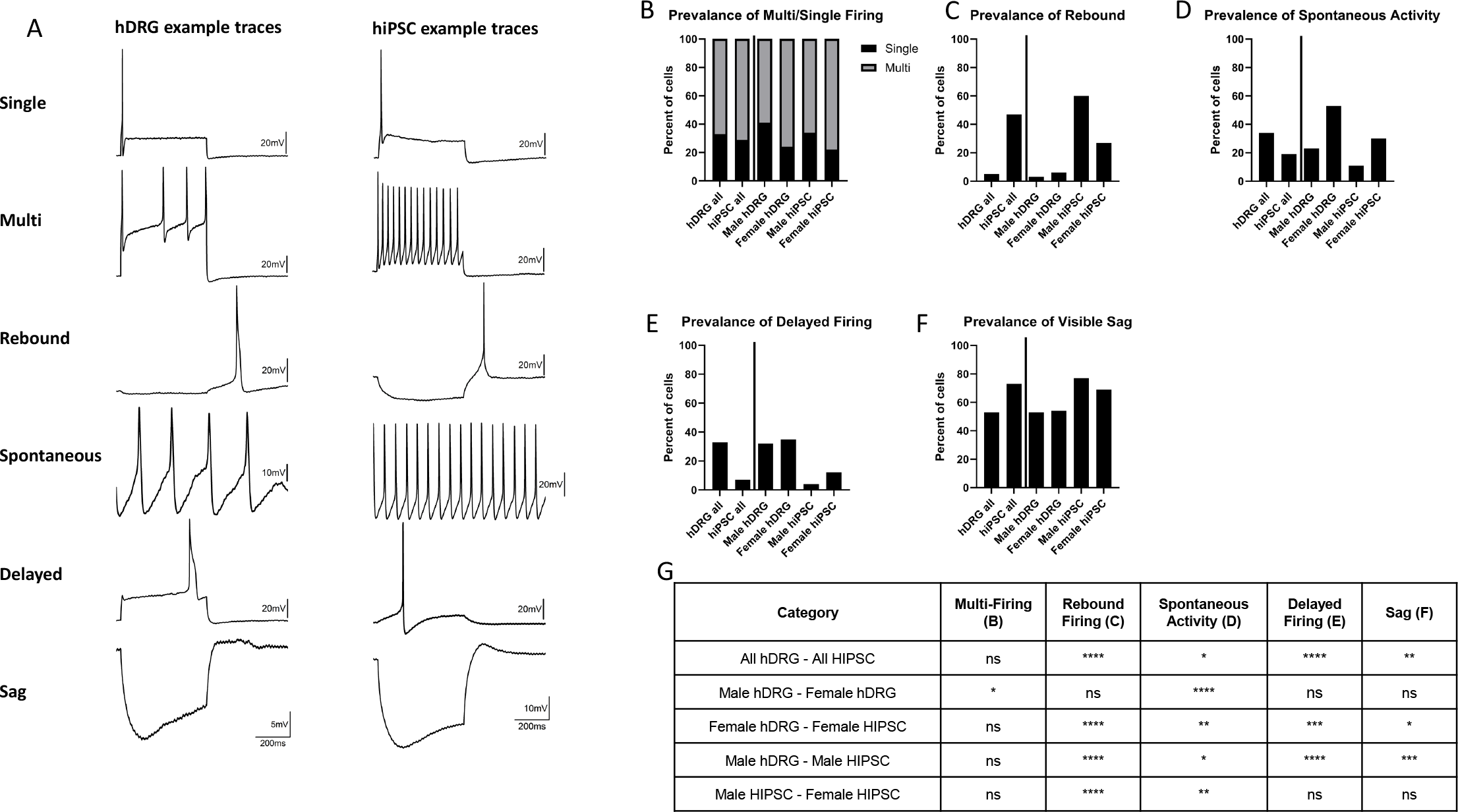
Firing Phenotypes. **(A)** Example traces of phenotypes observed in hDRG (n=108) and hiPSC-SNs (n=83) and percent of cells the firing pattern was observed including **(B)** prevalence of multi-versus single firing cells, **(C)** prevalence of rebound firing, **(D)** prevalence of spontaneous activity, **(E)** prevalence of delayed firing, and **(F)** prevalence of visible sag (more than 10% sag ratio). **(G)** Statistical comparison between groups of the comparisons using Fisher’s Exact Test. *p=.01, **p<.01, ***p<.001, ****p<.0001.

### Waveform characteristics of hDRG-N compared to hiPSC-derived SNs

#### AP amplitude

Differences in AP waveform characteristics were analyzed for all cells. There were significant differences in many AP waveform properties between hDRG-N and hiPSC-SN. The AP peak was higher for hDRG-N at 52.65 ± 1.343 mV versus 39.80 ± 1.273 for hiPSC-SN, p < 0.0001 (Figure 5A). The AP amplitude was similar between hDRG-N and hiPSC-SNs (78.23 ± 1.824 mV and 74.43 ± 1.325) (Figure 5B). The AP threshold measured using Method II[34], was significantly different between hDRG-N and hiPSC-SNs (−25.58 ± 0.9830 mV vs −34.43 ± 0.6729 mV, p <0.0001) (Figure 5C). The AP after hyperpolarization (AHP) was significantly different between hDRG-N (−30.49 ± 0.738 mV), and hiPSC-SNs (−23.59 ± 0.615 mV), p < 0.0001. Taken together these data show that while the AP amplitude is similar for hDRG-N and hiPSC-SNs, the hiPSC-SNs start the AP at a lower threshold, therefore the AP peak is lower for the hiPSC-SN.

**Figure 5.**
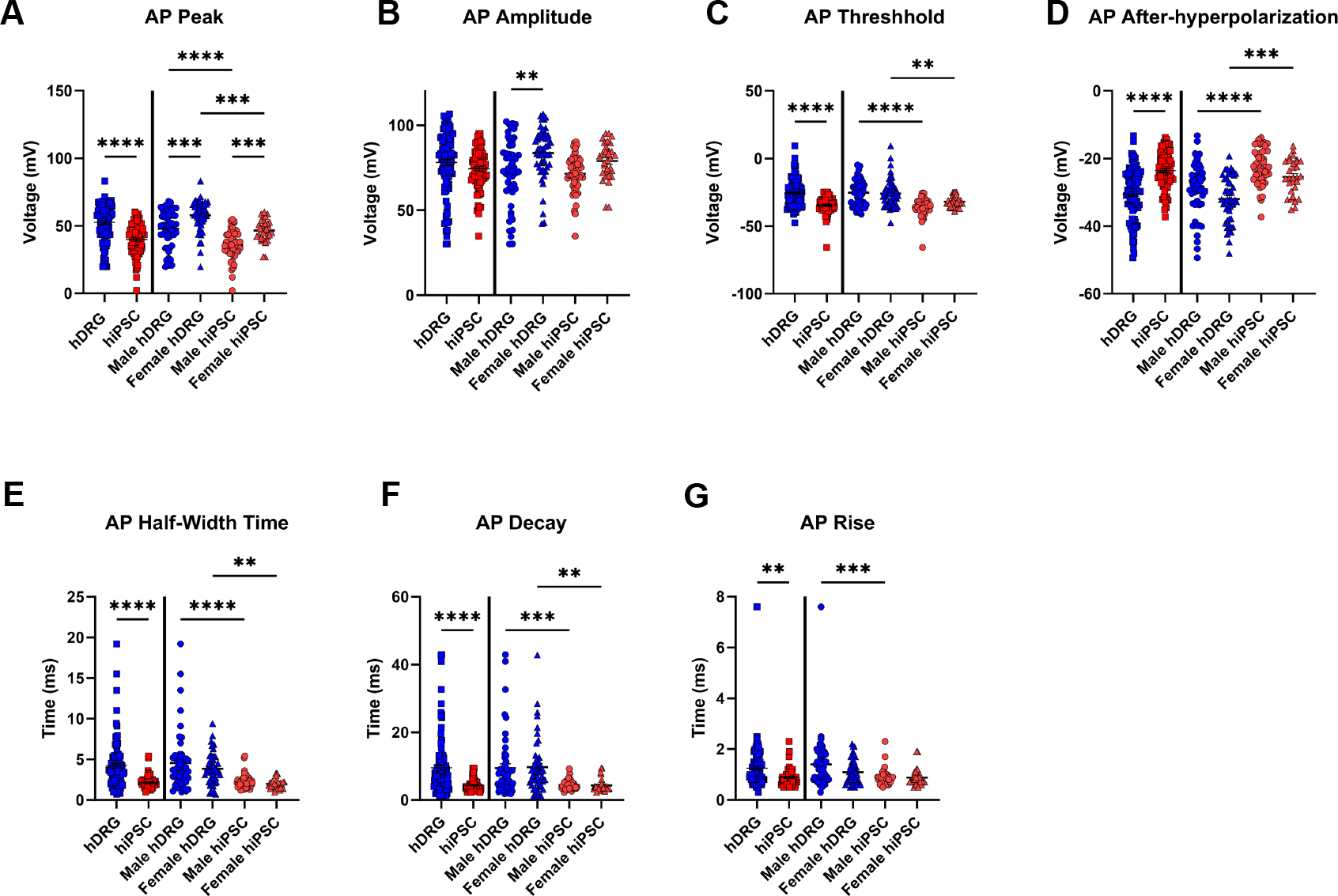
Action potential waveform characteristics. Differences in several action potential waveform characteristics were observed between hDRG cells (n=108) and hiPSC-SNs (n=82), **(A)** AP peak (mV), **(B)** AP amplitude (mV), **(C)** AP threshold (mV), **(D)** after-hyperpolarization **(E)** AP rise time (ms), **(F)** AP decay time (ms), **(G)** and AP half-width time (ms). Ordinary one-way ANOVA was used to determine statistical significance between cells from male hDRG (n=59), female hDRG (n=49), male hiPSC-SNs (n=50), and female hiPSC-SNs (n=33). Šídák’s multiple comparisons test confirmed differences in AP waveforms. *p=.01, **p<.01, ***p<.001, ****p<.0001. Error bars represent mean and SEM.

#### AP duration

AP duration was analyzed by measuring the AP rise time, decay time, and half-width. The AP rise time was significantly different between hDRG-N and hiPSC-SNs (1.251 ± 0.082 ms and 0.8846 ± 0.352 ms, p = 0.0010) (Figure 5E). The AP decay time was significantly longer in the hDRG-N at 9.541 ±0.8512 ms than hiPSC-SNs at 4.339 ± 0.1723 ms, p < 0.0001 (Figure 5F). The AP half-width was also significantly longer in hDRG-N (4.214 ± 2903 ms), than in hiPSC-SNs (2.137 ± 0.08499 ms), p < 0.0001 (Figure 5G). The range in all AP duration statistics was much bigger for hDRG-N than in hiPSC-SNs. AP rise range was 7.3 ms for hDRG-N and 1.8 ms for hiPSC-SN. The AP decay range was 41.7 ms for hDRG-N and 7.117 ms for hiPSC-SNs. The half-width range was 18.5 ms for hDRG-N and 4.4 ms for hiPSC-SNs. These data show that the AP duration is more variable for hDRG-N, and the APs tend to be longer when compared to hiPSC-SNs.

### Phase plot analysis

Phase plots for APs were analyzed to determine the prevalence of an AP shoulder. APs with a shoulder (Figure 6A) have a distinct pattern in their phase plots where the lower part of the phase plot displays a noticeable hump (Figure 6B, shaded region). APs with no shoulder (Figure 6C) do not display this hump (Figure 6D, shaded region). Nearly all hDRG-N, 96% (95/99) had an AP shoulder while only a small percentage of hiPSC-SNs had an AP shoulder, 14% (11/78), p < 0.0001 (Figure 6E). When averaged (Figure 6F), the phase plots showed that hDRG-N and hiPSC-SNs were more different from each other than the sex differences between the different phase plots within each group.

**Figure 6.**
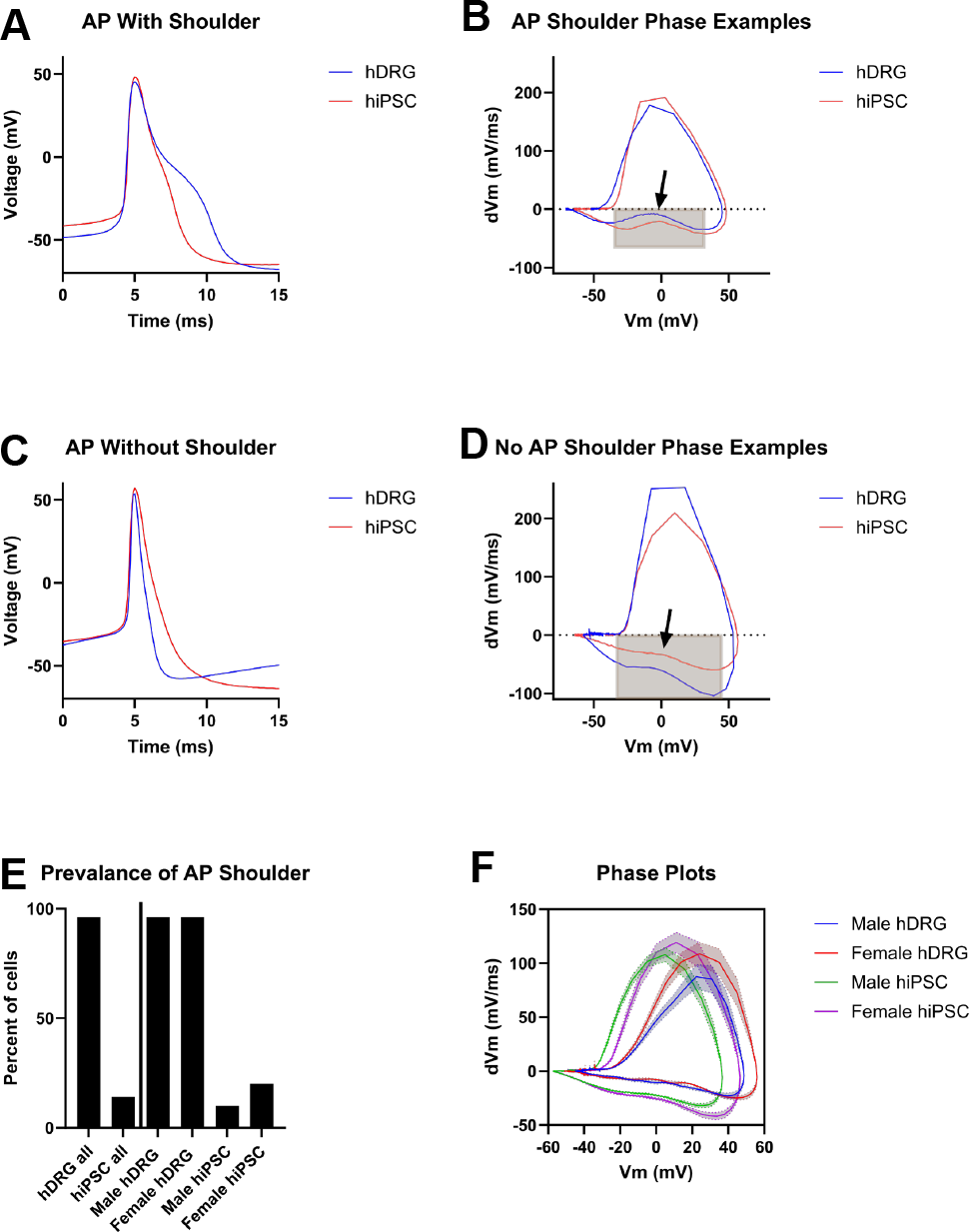
Phase Plot Characterization. **(A)** Representative action potentials with a shoulder. **(B)** Representative phase plots with a shoulder where the shaded area shows AP shoulders. **(C)** Representative action potentials without a shoulder, shaded area shows lack of AP shoulder. **(D)** Representative phase plots without a shoulder. **(E)** Prevalence of AP shoulder in all cell types. **(F)** Combined average of all hDRG and hiPSC-SNs phase plots, shaded area represents SEM.

### Correlations between electrophysiological properties

Pearson correlation matrices were used to determine correlations between the different electrophysiological properties of the cells. hDRG-N cells that had a high rheobase were less likely to be multi-firing or have a greater number of APs (Figure 7A). APs of high rheobase hDRG-N cells were of shorter duration measured by the half-width. The rheobase was also negatively correlated with the R_in_ in hDRG-N but positively correlated with the capacitance. In hDRG-N, R_in_ was also negatively correlated with the capacitance and the AHP. In hiPSC-SN some of these correlations were similar, rheobase was negatively correlated with Max APs and R_in_; however, there was little correlation between rheobase and half-width (Figure 7D). As in hDRG-N, R_in_ in hiPSC-SN was negatively correlated with capacitance and AHP, but only in female hiPSC-SN when the correlation plots were categorized by sex (Figure 7E, F).

**Figure 7:**
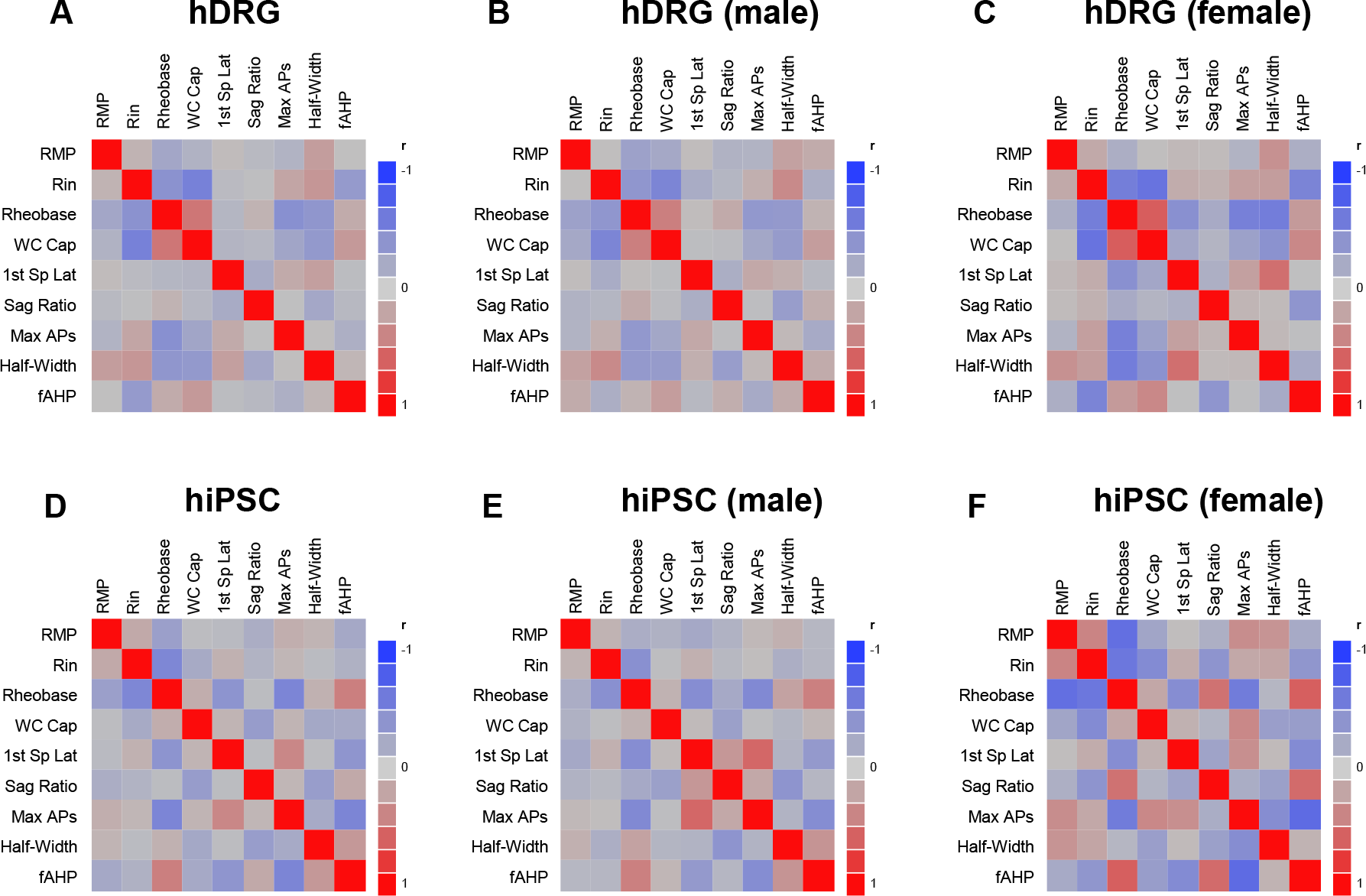
Pearson correlation matrices among hDRG and. hiPSC-SNs for RMP, Rin, rheobase, first-spike latency (1st Sp Lat), sag Ratio, and maximum count of action potentials (max APs), Half-width, and AHP for (**A)** hDRG pooled-sex, (**B)** hDRG male, (**C)** hDRG female cells, (**D)** hiPSC-SNs pooled-sex, (**E)** hiPSC-SNs male, and (**F)** hiPSC-SNs female cells. Diagonals show r=1.0

### Sex Differences in hDRG-N and hiPSC-SN

#### hDRG-N

In addition to comparing the differences between hDRG-N and hiPSC-SN, sex differences within each group were also compared. hDRG-N showed no difference in cell diameter, RMP, rheobase, R_in_, capacitance, or FSL (Figure 2). There were no significant sex differences in hDRG-N excitability by f-I relationships and inactivation threshold (Figure 3). Some sex differences in hDRG-N were detected among firing patterns. There was a statistical sex difference in percentage of multi-firing hDRG-N, 59% (35/59) of male hDRG-N versus 75% (37/49) of female hDRG-N, p = 0.0154, cells were multi-firing, and in spontaneous activity, 22% (13/57) of male cells compared to 53% (18/34) of female cells had spontaneous activity, p < 0.0001. There was no sex difference in rebound, delayed firing, or visible sag (Figure 4). There were also sex differences in AP waveform properties of hDRG-N. Male hDRG-N had an AP peak of 47.88 ± 1.907 mV while female hDRG-N had an AP peak of 57.72 ± 1.615 mV, p = 0.0003. There was a slight sex difference in AP amplitude in hDRG-N (73.10 ± 2.734 mV for male and 83.67 ± 2.159 mV for female, p = 0.0034). There were no sex differences in AP threshold and AHP. There was also no sex difference in AP duration, measured by AP rise, AP decay, and AP half-width in hDRG-N (Figure 5). There was no difference in presence of AP shoulder between male and female hDRG-N (Figure 6).

#### hiPSC-SN

hiPSC-SN had a sex difference in RMP (−58.04 ± 1.022 mV in male cells and −53.06 ± 1.175 mV in female cells, p = 0.0214) and no sex differences in any other intrinsic properties, cell diameter, rheobase, R_in_, capacitance, sag ratio, and FSL (Figure 2). There was also no sex difference in f-I plots or inactivation threshold (Figure 3). There were two sex differences noted in firing patterns in hiPSC-SN, rebound firing was 60% (30/50) in male and 29% (9/33) in female cells (p < 0.0001), and in spontaneous activity, 11% (5/45) of male cells compared to 30% (9/30) of female cells (p = 0.0014). There were no significant sex differences in multi-firing cells, delayed firing, and cells with visible sag (Figure 4). There was only one sex difference in AP waveform properties, male hiPSC-SN had an AP peak of 35.57 ± 1.566 mV, and female hiPSC-SNs had an AP peak of 46.56 ± 1.503 mV, p = 0.0005 (Figure 5A). Other waveform properties showed no sex difference in hiPSC-SN (Figure 5). Only a slight sex difference in AP shoulder prevalence was detected between male, 10% (5/43) and female, 20% (6/24), hiPSC-SNs, p = 0.0477 (Figure 6).

## DISCUSSION

Several groups have made use of hDRG-N[1,8,12,20–23,27,35,36,39,43] and hiPSC-SNs [4,28,30,32] in studies of neuronal excitability using patch-clamp electrophysiology. This is the first analysis of electrophysiological recordings obtained from the same lab of both cell types. A comparison of these tools is vital for understanding the advantages and drawbacks of each. In this study, the electrophysiological properties for hDRG-N and hiPSC-SNs [41] were compared. As major points, the results of this study revealed hDRG-N in general have a larger range of values for intrinsic properties such as cell diameter, rheobase, capacitance, and first spike latency, while hiPSC-SNs had a small range for these properties and were less variable. hiPSC-SNs tended to have a lower rheobase and started multi-firing at a lower current above rheobase than hDRG-N. Phenotypic differences in firing patterns were seen with hiPSC-SNs having more rebound firing and a lower frequency of delayed firing than hDRG-N. hDRG-N tended to have a higher AP threshold however the AP amplitude was similar to hiPSC-SNs. hDRG-N had a wider range of AP durations measured by AP rise, half-width, and AP decay, while hiPSC-SNs had more consistent AP duration.

When looking at AP phase plots we found that 96% of hDRG-N had an AP shoulder, consistent with findings of other groups[7], while only 14% of hiPSC-SNs showed an AP shoulder. Previous work has found that the presence of an AP shoulder and longer half-width of hDRG-N is contingent on presence of functional Nav1.8 in the membrane[3,12]. In our previous work we have shown that these hiPSC-SNs express Nav1.7, Nav1.8, and Nav1.9 mRNA like hDRG-N[38]. However, the short half-width and lack of an AP shoulder in hiPSC-SN suggest the protein may not be functional or that is it not localized to the membrane of hiPSC-SNs. More work is needed to understand the role of Nav1.8 and other ion channels in hiPSC-SNs and how they contribute to differences in the AP waveform properties compared to hDRG-N.

Sex differences are a major area of interest in chronic pain research. Molecular differences have been identified in sensory neurons from male and female donors using molecular techniques such as transcriptomics and translatomics[31,37]. A key difference between male and female nociceptors was found in prostaglandin signaling, which relates to inflammation[37]. While this may differentially influence the response of nociceptors from male and female neurons to sensitizers, it perhaps will not change their intrinsic electrophysiological properties that relies more on ion channel expression. Our results reveal greater potential differences between hiPSC-SNs and hDRG-N versus potential sex differences within each group. We only detected a few small sex differences. In hDRG-N we only saw sex differences in percentage of multi-firing cells, percentage of spontaneous activity, AP amplitude, and AP peak. In hiPSC-SN we saw sex differences only in RMP, rebound firing, delayed firing, AP peak, and AP shoulder prevalence. Most of intrinsic, phenotypic, and AP waveform properties were significantly different between hDRG-N and hiPSC-SN showing that sex differences are less prominent than the differences between cell types.

Correlations between intrinsic and waveform properties demonstrate several positive and negative correlations between properties. Taken together these show that both hDRG-N and hiPSC-SNs could be useful tools in understanding neuronal excitability though for different reasons. hDRG-N show the true dynamic range of neuronal cells within human dorsal root ganglion tissue while hiPSC-SNs are more consistent and could be more useful for detecting differences between different treatment paradigms with a smaller sample set.

hDRG-N have been used in several studies to look at effects of various compounds in potentiating and attenuating neuronal excitability[2]. Because hDRG-N contain a variety of neuronal subtypes such as nociceptors and mechanoreceptors, data analysis requires a large dataset as well as separating neurons by subtype based on firing patterns such as multi versus single firing neurons, rheobase, and AP waveform properties[6,16,43]. hDRG-N cultures also contain other cell types such as satellite glia, therefore some effects of the compound being tested may be due to interactions with the other cell types. This makes hDRG-N advantageous for studies looking at the properties of the DRG holistically rather than as a single cell type. However, it also makes hDRG-N data analysis difficult, requiring researchers to separate cells by firing properties without knowing the neuronal cell type[43] or the need for more advanced techniques such as patch-seq[29], which has yet to be published in human sensory neurons.

On the other hand, hiPSC-SNs have been introduced as a valuable tool for looking at effects on a single cell type. hiPSC-SNs have been generated by several different groups using a variety of methods[4,9,14,17–19,24–26,28,32,44]. They are clonal and have more consistent properties such as rheobase and AP duration, making them a valuable tool in screening for compounds that affect neuronal excitability. In this study hiPSC-SNs from Anatomic Corp. derived from human peripheral blood mononuclear cells were utilized for comparison as these cells are currently widely used by several pain labs in the field[41]. The differences between methods used to generate hiPSC-SNs could lead to biological differences; therefore, using hiPSC-SN from a single verified source allows for comparison between samples. The consistent firing properties within each hiPSC-SN culture allows researchers to screen compounds more rapidly compared to hDRG-N. However, hiPSC-SNs are dissimilar from hDRG-N in many ways. Since they do not have any support cells such as satellite glia and do not contain multiple neuronal subtypes, effects of compounds can only be determined as the effect on the hiPSC-SNs alone. While hiPSC-SNs express many of the mRNA and proteins that are expressed in hDRG-N[5,18,27,32,42,44], their expression patterns sometimes differ and this is likely what leads to key differences in firing patterns and AP waveform, such as the lack of an AP shoulder in hiPSC-SNs that is present in almost all hDRG-N.

Both hDRG-N and hiPSC-SNs are useful tools in pain research and each has advantages and potential challenges. hiPSC-SNs have a distinct advantage in that they can be derived from living pain patients and have the potential for use in personalized medicine in the future[19]. hDRG-N are usually derived from recently deceased patients or patients with loss of function below the lumbar region in complex surgeries[7,12,22,40]; therefore hDRG-N may lack the ability to be used in determining the efficacy of pain therapeutics for another specific patient. However, hDRG-N paint a more pragmatic picture of the behaviors of sensory neurons due to the diversity of cell populations present in hDRG-N cultures.

This study demonstrates that hiPSC-SNs have multiple key differences in electrophysiological properties when compared to hDRG-N derived nociceptors. Further work is required to establish whether these key differences limit the usefulness of hiPSC-SNs in pain research and personalized medicine. Additionally, the current study has only used hiPSC-SNs from people with no history of pain. Differences in neuronal excitability and mRNA expression have been reported in hDRG-N from patients with pain history versus patients with no pain history[5,13,15,20–22,43]. Future work will need to be done to determine if the differences in hDRG-N are reproducible in hiPSC-SNs from pain patients.

## ACKNOWLEDGEMENTS

We are grateful for the donors and their families and humbled to have the opportunity to do this research. Thanks to the organ donation and transplant team, who have supported and facilitated this project in numerous ways. We would like to acknowledge Anatomic Corp. for providing the hiPSCs.

